# A *Caenorhabditis elegans nck-1* and filamentous actin-regulating protein pathway mediates a key cellular defense against bacterial pore-forming proteins

**DOI:** 10.1101/2022.06.10.495598

**Authors:** Anand Sitaram, Yunqiang Yin, Tammy Zamaitis, Bo Zhang, Raffi V. Aroian

**Affiliations:** Program in Molecular Medicine, University of Massachusetts Chan Medical School, Worcester, Massachusetts, USA; Department of Neurology, Boston Children’s Hospital, Harvard Medical School, Boston, Massachusetts, USA; Biostatistics and Research Design Center, Institutional Centers for Clinical and Translational Research, Boston Children’s Hospital, Harvard Medical School, Boston, Massachusetts, USA

## Abstract

Pore-forming proteins (PFPs) comprise the largest single class of bacterial protein virulence factors and are expressed by many human and animal bacterial pathogens. Cells that are attacked by these virulence factors activate epithelial intrinsic cellular defenses (or INCEDs) to prevent the attendant cellular damage, cellular dysfunction, osmotic lysis, and organismal death. Several conserved PFP INCEDs have been identified using the nematode *Caenorhabditis elegans* and the nematicidal PFP Cry5B, including mitogen-activated protein kinase (MAPK) signaling pathways. Here we demonstrate that the gene *nck-1*, which has homologs from *Drosophila* to humans and links cell signaling with localized F-actin polymerization, is required for INCED against small-pore PFPs in *C. elegans*. Reduction/loss of *nck-1* function results in *C. elegans* hypersensitivity to PFP attack, a hallmark of a gene required for INCEDs against PFPs. This requirement for *nck-1*-mediated INCED functions cell-autonomously in the intestine and is specific to PFPs but not other tested stresses. Genetic interaction experiments indicate that *nck-1*-mediated INCED against PFP attack is independent of the major MAPK PFP INCED pathways. Proteomics and cell biological and genetic studies further indicate that *nck-1* functions with F-actin cytoskeleton modifying genes like arp2/3, *erm-1*, and *dbn-1* and that *nck-1/arp2/3* promote pore repair at the membrane surface and protect against PFP attack independent of p38 MAPK. Consistent with these findings, PFP attack causes significant changes in the amount of actin cytoskeletal proteins and in total amounts of F-actin in the target tissue, the intestine. *nck-1* mutant animals appear to have lower F-actin levels than wild-type *C. elegans*. Studies on *nck-1* and other F-actin regulating proteins have uncovered a new and important role of this pathway and the actin cytoskeleton in PFP INCED and protecting an intestinal epithelium *in vivo* against PFP attack.

**Author Summary:** The mechanism of action for a significant number of bacterial protein toxins is the formation of pores in the membrane of target cells. Host cells contain programmed defenses against such attacks. Here we use the model system of the roundworm *Caenorhabditis elegans* and crystal proteins produced by *Bacillus thuringiensis* to demonstrate a new defense pathway mediated by the activity of the NCK-1 protein. Consistent with its known function in mammalian cells, the NCK-1-mediated defense involves various actin-interacting proteins and affects the kinetics of pore repair. This pathway is novel in its independence from MAPK signaling. Furthermore, the NCK-1 activity is found to be generally needed for defense against multiple pore-forming proteins and yet is specific to defense against such proteins but not against other environmental stressors.

## Introduction

The largest class of bacterial protein virulence factors is pore-forming proteins (PFPs) [1–4]. Produced by many major human pathogens, these are critical virulence factors; many strains that have been deleted for their PFP genes become avirulent or significantly less virulent in lab animals. It is therefore critical to better understand how host cells respond to attack by these toxins. Typically, toxin monomers are secreted or externalized by bacteria, activated by cleavage by host proteases, and oligomerize before being inserted into the plasma membrane and forming pores. PFPs can generally be split into large pore-formers (∼30 nm diameter) and small pore-formers (∼2 nm diameter). The host range and cellular tropism can be determined by expression of host factors at the membrane that serve as receptors for the toxin. Receptors for different toxins include cholesterol, other lipids, and proteins. Cells that are perforated by toxins are thought to die by osmotic lysis, necrosis, programmed cell death, or dysfunction caused by ion dysregulation. [5].

Cells can mitigate this damage, however, by activating *in*trinsic *ce*llular *d*efenses or INCED [6–12]. The nematode *Caenorhabditis elegans* and exposure to PFPs (*e*.*g*., Cry5B PFP; [13]) made by the soil bacterium *Bacillus thuringiensis* have proven to be an invaluable model for discovering and studying INCED pathways by which cells defend against PFPs. These INCEDs include activation and up-regulation of p38 and JNK-like mitogen-activated protein kinase (MAPK) signaling pathways [13,14], the unfolded protein response [9], the hypoxia response [6], endocytosis and membrane shedding [15], and autophagy [7]. Investigation in animal cell culture systems has shown that all of these PFP INCED response processes discovered and/or described in *C. elegans* are evolutionarily conserved [9,13,14,16].

We previously reported on a *C. elegans* genome-wide RNAi screen for genes required for INCED against PFPs, in which >16,000 RNAi knock-downs were performed followed by selection and repeat testing at a single dose of Cry5B PFP [13]. This screen yielded 106 *hpo* genes (<u>h</u>ypersensitive to <u>po</u>re-forming toxin). One of the conserved hits included *nck-1*, the *C. elegans* homolog of mammalian Nck that is involved in linking tyrosine phosphorylation with localized F-actin polymerization by recruiting several effectors to the plasma membrane, including N-WASP and the Arp2/3 complex [17]. *nck-1* is known to be involved in *C. elegans* axonal guidance, excretory cell and distal tip cell migration, male mating, vulval development, dauer formation, and susceptibility to Orsay virus [18–22]. Here we study the role of *nck-1* in PFP INCED, uncovering a key, important, and previously unknown role *in vivo* between PFP INCED, *nck-1* and other F-actin modulating genes, and the filamentous (F-) actin cytoskeleton.

## Results

To follow up on the genome-wide RNAi screen for PFP INCED genes [13], we selected a few *hpo* genes for dose-dependent Cry5B PFP response assays using RNAi and also looked at the corresponding response of available genetic mutants associated with these *hpo* genes. From these preliminary follow up studies, the *hpo* gene *nck-1* emerged as a gene of interest. Wild-type N2 *C. elegans* larvae grown on *E. coli* expressing RNAi sequences directed against *nck-1* were subjected to dose-response mortality assays using purified Cry5B (Figure 1A). *nck-1(RNAi) C. elegans* hermaphrodites were hypersensitive to Cry5B relative to *empty-vector(RNAi) C. elegans* hermaphrodites over a wide range of Cry5B PFP. In these and other experiments below, RNAi of the mitogen-activated protein kinase kinase (MAPKK) *sek-1*, knockdown of which is known to give a high level of hypersensitivity to Cry5B PFP [14,15], was included as a positive control. These experiments confirm *nck-1* is required for normal PFP INCED over a wide range of PFP doses.

**Figure 1.**
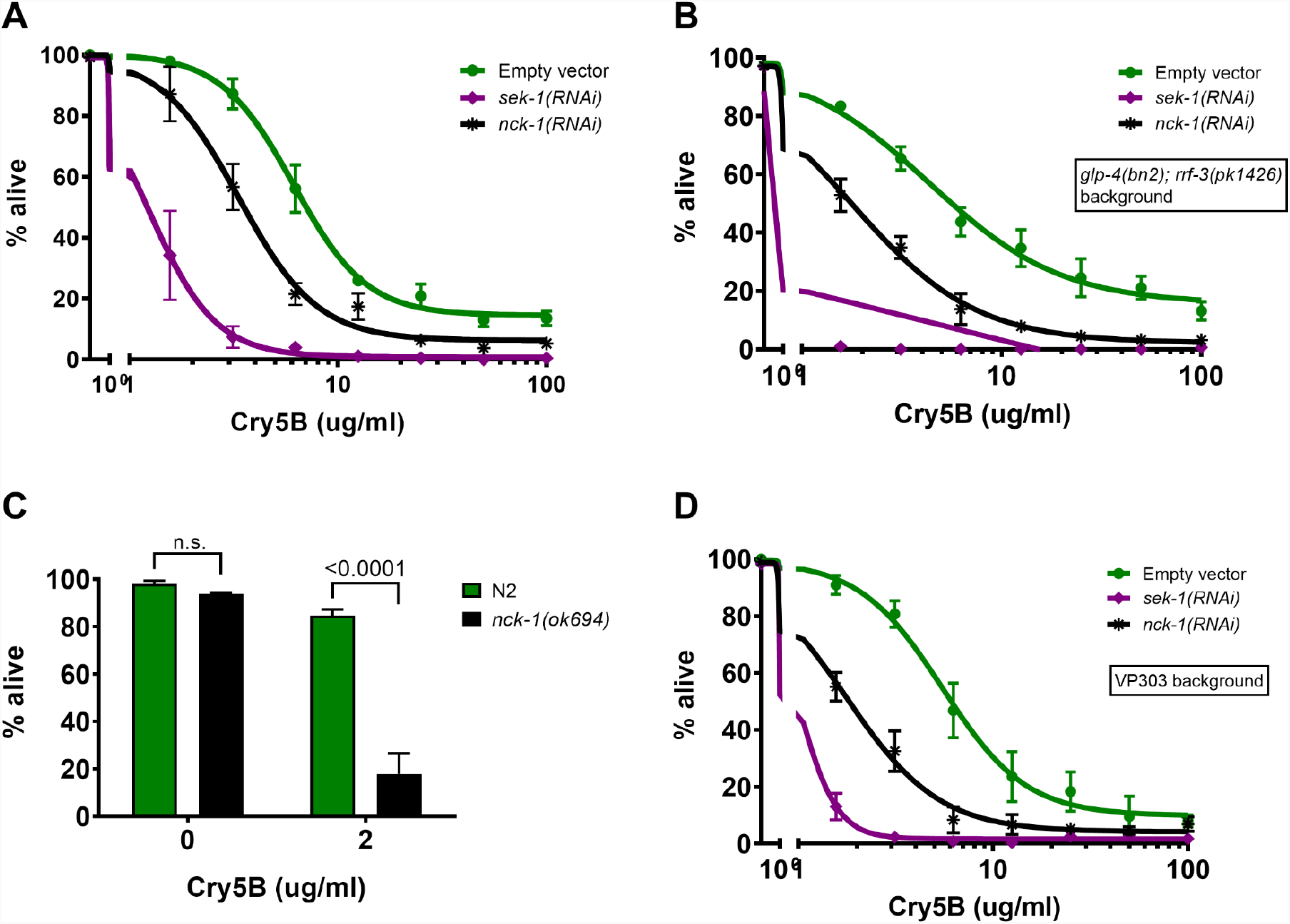
*nck-1* loss sensitizes *C. elegans* to Cry5B. (A) Wild-type N2, (B) *glp-4(bn2); rrf-3(pk1426)*, or (D) VP303 (intestine-specific RNAi) *C. elegans* were grown from the L1 to L4 stage on *E. coli* expressing the indicated RNAi, then transferred to liquid culture with purified Cry5B for 6 days, then assayed for survival. (C) Wild-type N2 or *nck-1(ok694)* null L4s were grown on OP50 *E. coli* until the L4 stage before being subjected to the same experiment. All data show the mean of three experiments. Error bars here and in other figures represent the standard error of the mean) (SEM).

These experiments have a time course of 6 days, and the presence of progeny born during that time can obscure the results, so the thymidine analog 5-fluoro-2’-deoxyuridine (FUdR) was included in the media in Figure 1A to prevent progeny production. To ensure that the presence of FUdR itself did not contribute to the Cry5B hypersensitivity of *nck-1*-deficient worms, the experiment was repeated using the *glp-4(bn2); rrf-3(pk1426)* strain, which is hypersensitive to RNAi (due to the *rrf-3* mutation) as well as sterile at the assay temperature of 25°C (due to the *glp-4* mutation), obviating the need for FUdR. In this background, knockdown of *nck-1* also led to hypersensitivity to Cry5B as measured by mortality (Figure 1B). The outcrossed null mutant *nck-1(ok694)* was viable and fertile, but larvae that were synchronized by standard hypochlorite treatment nevertheless developed at variable speeds (data not shown). Therefore L4 larvae were visually identified and handpicked for experiments using the mutant strain. As with wild-type worms knocked down for *nck-1* by RNAi, *nck-1(ok694)* mutants were also hypersensitive to Cry5B compared to wild-type worms, and the effect was more penetrant than in the RNAi experiment (Figure 1C). Because the intestinal epithelium is the tissue directly targeted by the PFP Cry5B [23–25], we also tested whether *nck-1* activity is required in the intestinal epithelium itself. Knockdown of *nck-1* in the VP303 strain—which preferentially carries out RNAi in the intestine—also led to hypersensitivity to Cry5B (Figure 1D), indicating that *nck-1* is required for a cell-autonomous defense process for protection against PFPs.

The p38 MAPK pathway, for which *sek-1* is the MAPKK, is clearly a central player in cellular defenses against PFPs ([5,13,26]. We next tested whether or not *nck-1*’s role in PFP defenses was part of the p38 MAPK PFP defense pathway. First we examined the MAPK-dependent activation of the *xbp-1-*dependent arm of the unfolded protein response (UPR) in response to Cry5B PFP. The *hsp-4::gfp* worm strain carries a GFP reporter for this response, increasing its fluorescence in response to classic UPR inducers such as tunicamycin or heat shock, but also after worms are fed Cry5B-expressing *E. coli* for 8 hours [9]. The latter, PFP-induced UPR response is known to be dependent upon MAPK signaling, as it was previously shown to be blocked in animals that carry a secondary mutation in *pmk-1*, which encodes the *C. elegans* p38 homolog that is a direct target of SEK-1 kinase activity. Here we similarly found that *hsp-4::gfp* worms that lacked MAPK signaling due to knockdown of *sek-1* showed no increase in GFP signal in response to an 8-hour *E. coli-*Cry5B feeding exposure (Figure 2A). In contrast, the knockdown of *nck-1* did not block Cry5B-induced activation of the UPR. Worms knocked down for *sek-1* or *nck-1* still showed increased GFP signaling following an 8-hour 30°C heat shock exposure, demonstrating that the *hsp-4::gfp* response was still intact in those worms (data not shown). As a control, worms with a knockdown for *xbp-1* showed a loss of even basal levels of reporter activity in response to either stressor (Figure 2A and data not shown). These results suggested a distinction between a *sek-1*-mediated and *nck-1*-mediated PFP response.

**Figure 2.**
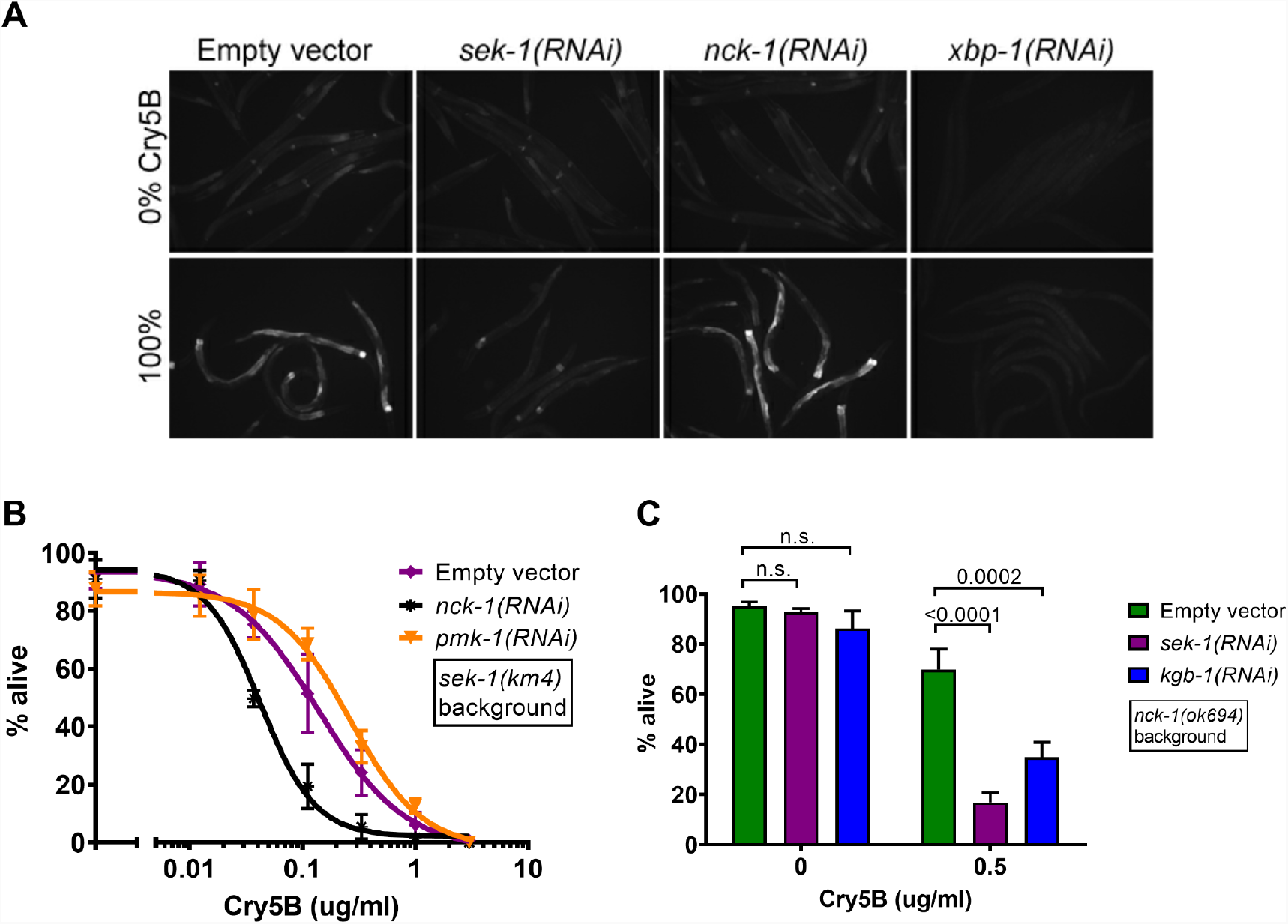
*nck-1* works independently of MAPK signaling. (A) *hsp-4::gfp C. elegans* were grown to the L4 stage on the indicated RNAi bacteria and transferred to *E. coli* plates expressing empty vector or Cry5B for 8 hours, then subjected to fluorescence microscopy. (B) *sek-1(km4)* p38 MAPKK(-) mutant *C. elegans* were grown on the indicated RNAi bacteria to the L4 stage and then transferred to liquid culture with varying amounts of purified Cry5B and incubated for 6 days before being scored for survival. *pmk-1* is p38 MAPK. (C) The reverse experiment to the one in (B), with *nck-1(ok694)* mutant animals and RNAi of MAPK pathway genes at a single concentration of Cry5B. (B) and (C) show the average of three independent experiments and error bars show SEM.

We therefore next directly tested for a genetic interaction between MAPK signaling and *nck-1* in toxin defense. *sek-1(km4)* p38 MAPKK mutant hermaphrodites were subjected to additional knock down of either *nck-1* or *pmk-1*. As expected, *pmk-1* knockdown did not increase the PFP sensitivity of hermaphrodites that were already null for the upstream activator *sek-1* (Figure 2B). In contrast, *nck-1* knockdown did further sensitize *sek-1(km4)* null hermaphrodites to Cry5B. We also performed this experiment in reverse, knocking down either *sek-1* or *kgb-1* (part of the JNK-like MAPK signaling pathway in *C. elegans* that has also been critically implicated in PFP defense [13,14] in *nck-1(ok694)* null mutants. Knockdown of either MAPK pathway further sensitized *nck-1(ok694)* worms to Cry5B (Figure 2C). Taken together, these data demonstrate that *nck-1* functions independently of the p38 and JNK-like MAPK signaling pathways for Cry5B PFP defense.

We next characterized the specificity of the *nck-1*-mediated PFP response. Namely, is the hypersensitivity of *nck-1* reduction/loss-of-function to Cry5B PFP due to a specific defect in Cry5B PFP responses or to a more generalized inability to protect against even non-PFP stressors that attack the health of the nematode? We therefore next tested the ability of *nck-1*-deficient animals to withstand other types of stress. Worms that experienced *nck-1* RNAi in the intestine alone (the VP303 strain) showed no change in their sensitivity to a prolonged 35°C heat stress (Figure 3A). The intestinal knockdown of *nck-1* also did not sensitize worms to the heavy metal copper sulfate (Figure 3B). Global knockdown of *nck-1* in N2 did not lead to a change in sensitivity to osmotic stress (Figure 3C) or oxidative stress either (Figure 3D), as assayed by exposure to increasing concentrations of NaCl or H_2_O_2_, respectively. Since the tested stresses mentioned to this point are all environmental rather than pathogenic, we also tested the requirement for *nck-1* in defense against the pathogenic food source *Pseudomonas aeruginosa* (PA14), which produces several toxic factors but is not known to produce a PFP. Knockdown of *nck-1* did not sensitize the worms to PA14 (Figure 3E). Consistent with previous reports [27], *sek-1* knockdown did sensitize worms to PA14 feeding. In all specificity experiments, some empty vector-treated and *nck-1(RNAi)* worms were tested in parallel to confirm that the RNAi was effective (*i*.*e*., that the *nck-1(RNAi)* worms were hypersensitive to feeding on *E. coli* expressing Cry5B) (data not shown). Taken together, the results suggest that, whereas some pathways, *e*.*g*., p38 MAPK signaling, are known to be utilized as a defense against PFPs as well as other stresses [27–29], the use of *nck-1* to protect cells from PFPs appears to be more narrow.

**Figure 3.**
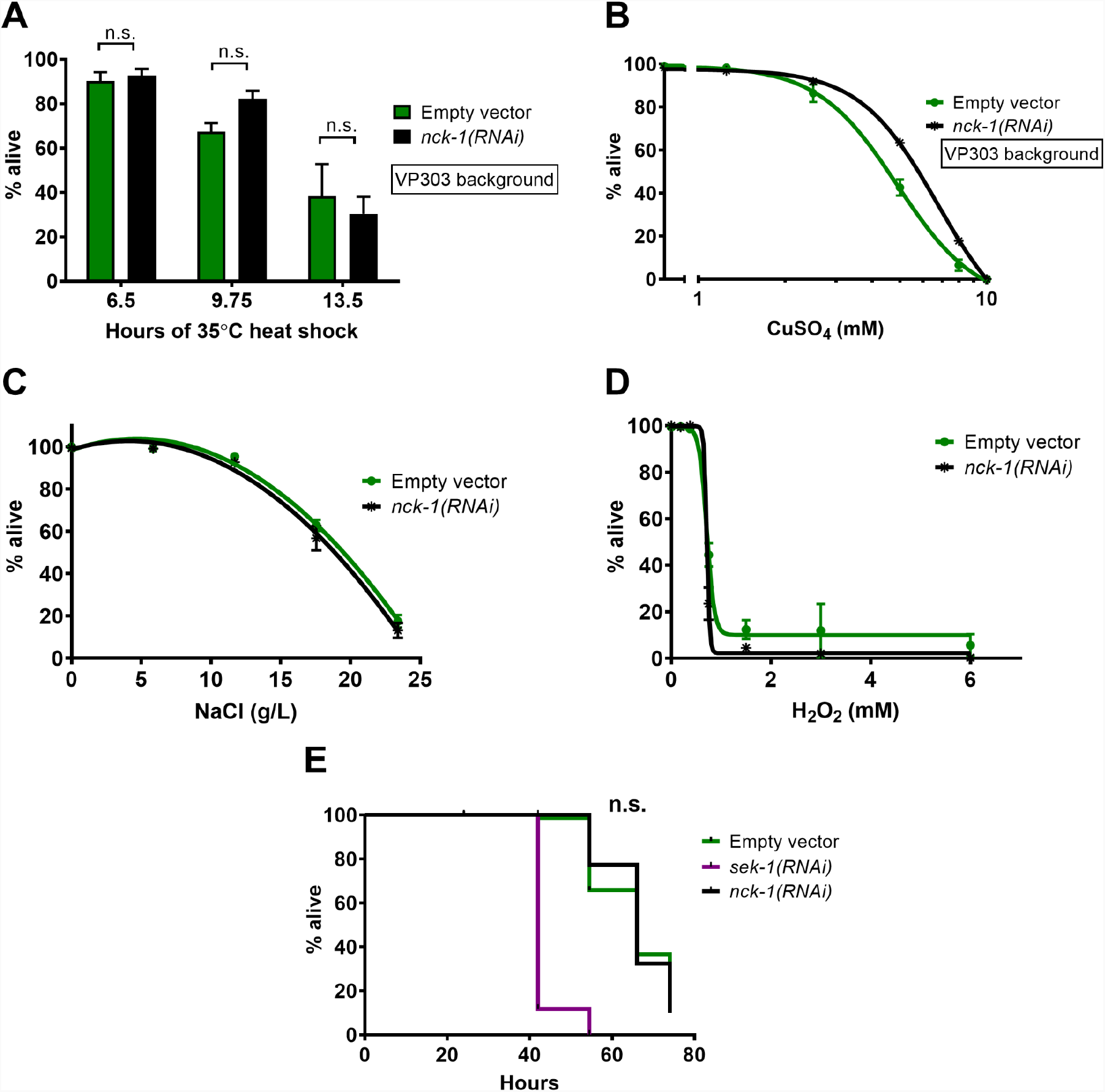
*nck-1* loss does not affect sensitivity to other stressors. (A) VP303 (intestinal-specific RNAi) *C. elegans* L4 animals grown on RNAi bacteria were incubated at 35°C, with survival determined at each indicated time-point. (B) VP303 L4 animals grown on the indicated RNAi bacteria were transferred to liquid culture with varying amounts of CuSO_4_ for 6 days at 25°C. (C and D) Wild-type N2 *C. elegans* grown to the L4 stage on the indicated RNAi were transferred to liquid culture with varying amounts of (C) NaCl for 6 days or (D) H_2_O_2_ for 4 hours, before survival was measured. (E) Wild-type *C. elegans* grown to the L4 stage on the indicated RNAi were transferred to plates spread with *Pseudomonas aeruginosa* PA14 and survival was followed over the indicated time. In (A)-(D) the data shown are the mean of three independent experiments, with error bars representing the SEM. The data in (E) are representative of three trials. Logrank analysis shows no significant difference in survival between empty vector and *nck-1(RNAi)* animals.

We then investigated whether or not *nck-1* protects *C. elegans* against PFPs other than Cry5B. For this, we tested the requirement for *nck-1* in defense against another member of the *Bacillus thuringiensis* crystal protein family of PFPs, App6A (formerly called Cry6A; [30]). Although Cry5B and App6A are both *B. thuringiensis* proteins that form pores, the two PFPs are structurally distinct and use different receptors, and *C. elegans* mutants that are resistant to Cry5B remain sensitive to App6A [13,31–35]. Hence, App6A represents an independent PFP for testing with *nck-1* animals. We found that *nck-1(ok694)* null mutants were visually hypersensitive to feeding on *E. coli* expressing recombinant App6A, compared to wild-type worms (Figure 4A).

**Figure 4.**
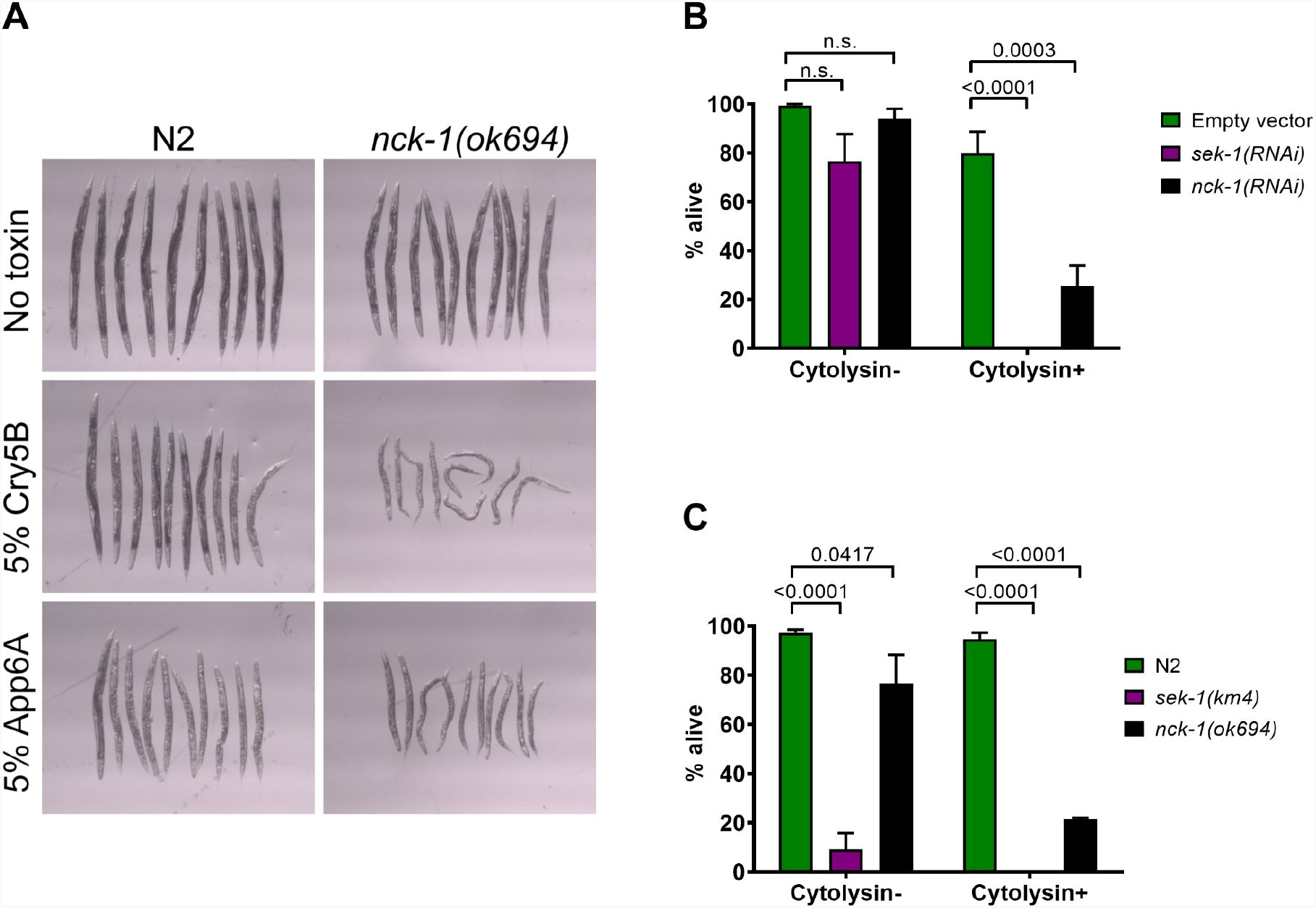
*nck-1* loss sensitizes *C. elegans* to multiple PFPs. (A) Wild-type N2 or *nck-1(ok694)* null *C. elegans* L4s were transferred to plates spread with the indicated percentage of *E. coli* expressing Cry5B or App6A (formerly Cry6A). Plates were incubated two days at 20°C and then worms were transferred to spot wells and photographed. (B) Wild-type N2 *C. elegans* grown to the L4 stage on the indicated RNAi bacteria were transferred to plates spread with *Vibrio cholerae* that do or do not express the PFP cytolysin, and survival was measured after 48 hours. (C) The same experiment as in (B) but using wild-type N2 or mutant *C. elegans* L4 animals. The data in (A) are representative of three trials. In (B) and (C), the data represent the mean of three trials and SEM.

We also tested the sensitivity of *C. elegans* to *Vibrio cholerae* cytolysin, a PFP that is active against both mammalian cells and *C. elegans [6,36]. C. elegans* with an *nck-1* deficiency due to either RNAi knockdown (Figure 4B) or genetic mutation (Figure 4C) were much more sensitive to feeding on a cytolysin-positive strain of *V. cholerae* compared to wild-type worms. *C. elegans* with a *sek-1* deficiency were also much more sensitive to feeding on a cytolysin-positive strain of *V. cholerae* compared to wild-type worms (Figure 4B). The increased sensitivity of *nck-1-*deficient animals compared to wild-type animals was not retained when fed *V. cholerae* lacking cytolysin, although *sek-1(km4)* null animals (but not *sek-1(RNAi)* reduction of function animals) were also hypersensitive to the cytolysin-minus strain of *V. cholerae*, consistent with the fact that *sek-1* is more generally involved in stress responses and that *V. cholerae* may produce other non-PFP virulence factors against *C. elegans* [37]. Taken together, these data demonstrate that *nck-1* shows specificity for defense against multiple PFPs but not the other environmental or pathogenic factors tested.

*C. elegans* intestinal cells that have pores inserted into their apical surface take steps to repair the integrity of the perforated membrane [15]. When worms are given an acute exposure to Cry5B PFP and then immediately fed the fluorescent dye propidium iodide, the ingested dye leaks from the intestinal lumen into the cytoplasm of the intestinal epithelial cells (Figure 5A; top panels). If, however, Cry5B PFP feeding and subsequent propidium iodide dye loading are separated by a ∼24-hr recovery period, the ingested dye is confined to the intestinal lumen, as the pores are repaired in the interim (Figure 5A; lower panels) [15]. Loss of RAB-11.1 activity due to partial RNAi knockdown weakens the cell’s ability to repair pores [15]. We asked whether or not NCK-1 was also involved in cellular pore repair. We found that whereas wild-type worms showed almost 100% repair during the ∼24-hr recovery period, *nck-1(ok694)* animals failed to repair dye-permeable pores, similar to *rab-11*.*1* control knockdowns (Figure 5B). The *nck-1(ok694)* mutant did not exhibit an endogenous intestinal permeability to propidium iodide in the absence of PFP exposure (data not shown).

**Figure 5.**
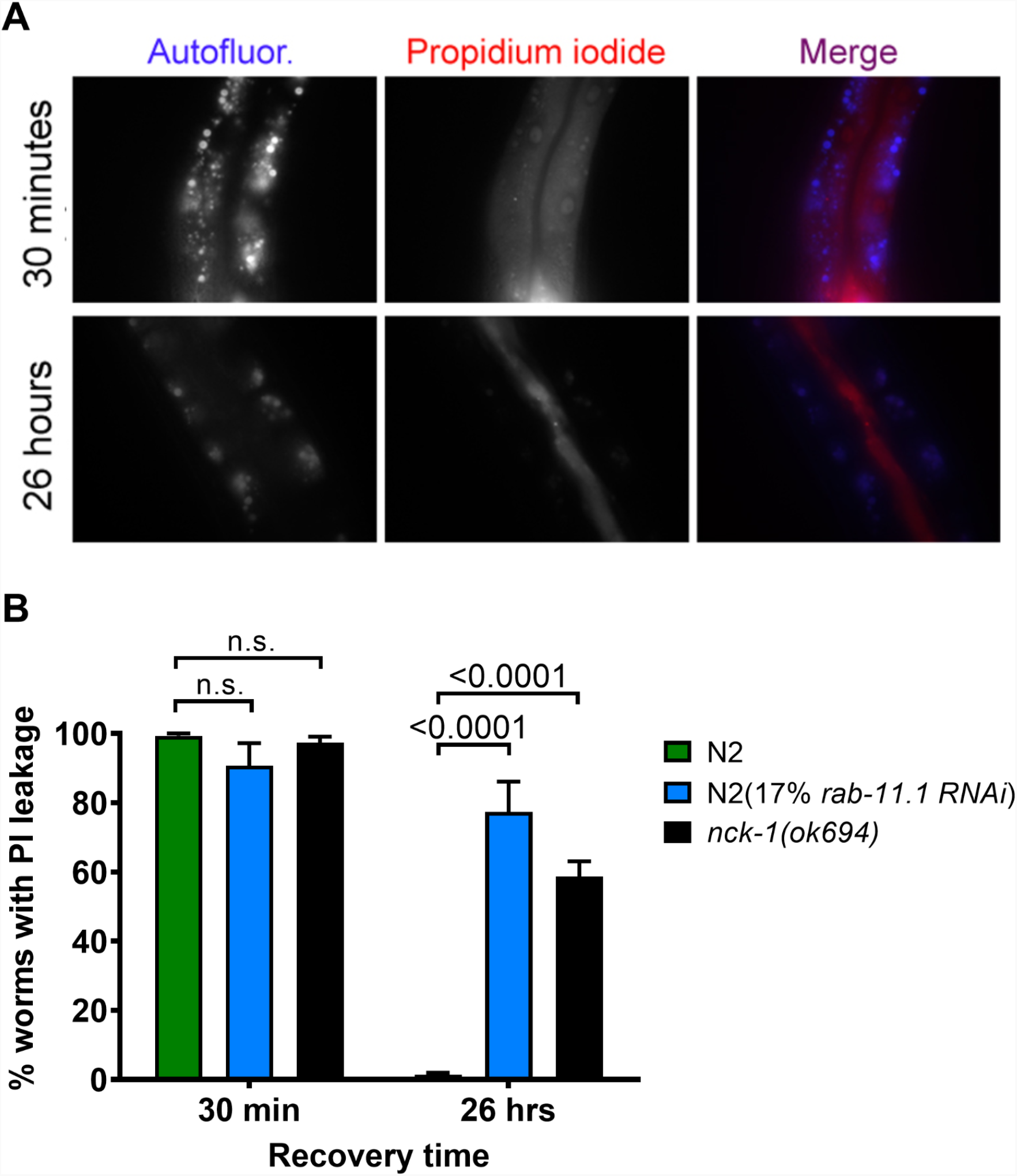
*nck-1* loss affects the ability of cells to repair PFP pores. (A) Wild-type *C. elegans* were grown to the L4 stage on the indicated RNAi bacteria and transferred to Cry5B plates for 1 hour. *C. elegans* were then either fed propidium iodide within 30 minutes or allowed to recover for 26 hours and then fed propidium iodide. Dye-loaded worms were imaged on slides. Autofluorescent gut granules serve as a surrogate marker indicating the position of the intestinal epithelial cells. (B) Quantitation of the same type of experiment as in (A) except the strains used were *nck-1(ok694)* null mutants (grown on standard OP50 *E. coli*) and N2 (grown on either OP50 or *rab-11*.*1* RNAi *E. coli* as indicated). “PI leakage” = animals with dye found in intestinal cell lumen. The data represent the mean and SEM of three independent experiments.

Following our finding that *nck-1* was required for efficient pore repair, we considered how it might function. In mammalian cells, the NCK-1 homolog Nck is an SH2/SH3 adaptor protein that recruits a large number of effector proteins to the plasma membrane following its own recruitment to phosphorylated tyrosine residues. In particular, Nck has been shown to be a potent activator of actin assembly, in part by effectively activating Wiskott-Aldrich Syndrome Protein (WASP)-family proteins so that they in turn activate the Arp2/3 complex, which initiates branched actin filament polymerization. The genome-wide RNAi and toxin hypersensitivity screen that identified *nck-1* as a *C. elegans* PFP defense gene also identified WASP-interacting protein *wip-1* as well as three subunits of the *C. elegans* Arp2/3 complex—*arx-3, arx-5*, and *arx-7—*as *hpo* (PFP defense) genes [13]. Based on these observations, we hypothesized that the *nck-1*-mediated response to PFPs might also function via the Arp2/3 complex.

We confirmed that Arp2/3 subunits are involved in PFP defense. Feeding of undiluted RNAi bacteria against *arx-3* or *arx-5* led to a significant growth defect in wild-type *C. elegans* (data not shown), so the RNAi bacteria were diluted 1:1 with bacteria carrying an empty vector. The level of knockdown afforded by this (50%) dilution was sufficient to qualitatively sensitize worms to Cry5B (Figure 6A). Wild-type worms knocked down for *arx-5* also show impaired pore repair (Figure 6B), a phenotype shared with *nck-1* mutants. We then looked for genetic interactions between *arx-5* and *nck-1*. Knockdown of *arx-5* in the *nck-1(ok694)* null background did not increase or decrease the sensitivity of the worms to Cry5B, whereas knockdown of *sek-1* in the *nck-1(ok694)* null background displayed increased sensitivity (Figure 6C; just as in the experiment in Figure 2C). Taken together, these results indicate that *nck-1* works in the same PFP defense pathway as the Arp2/3 complex but separate from the *sek-1* MAPK pathway, consistent with the relationship between Nck and Arp2/3 derived from mammalian studies.

**Figure 6.**
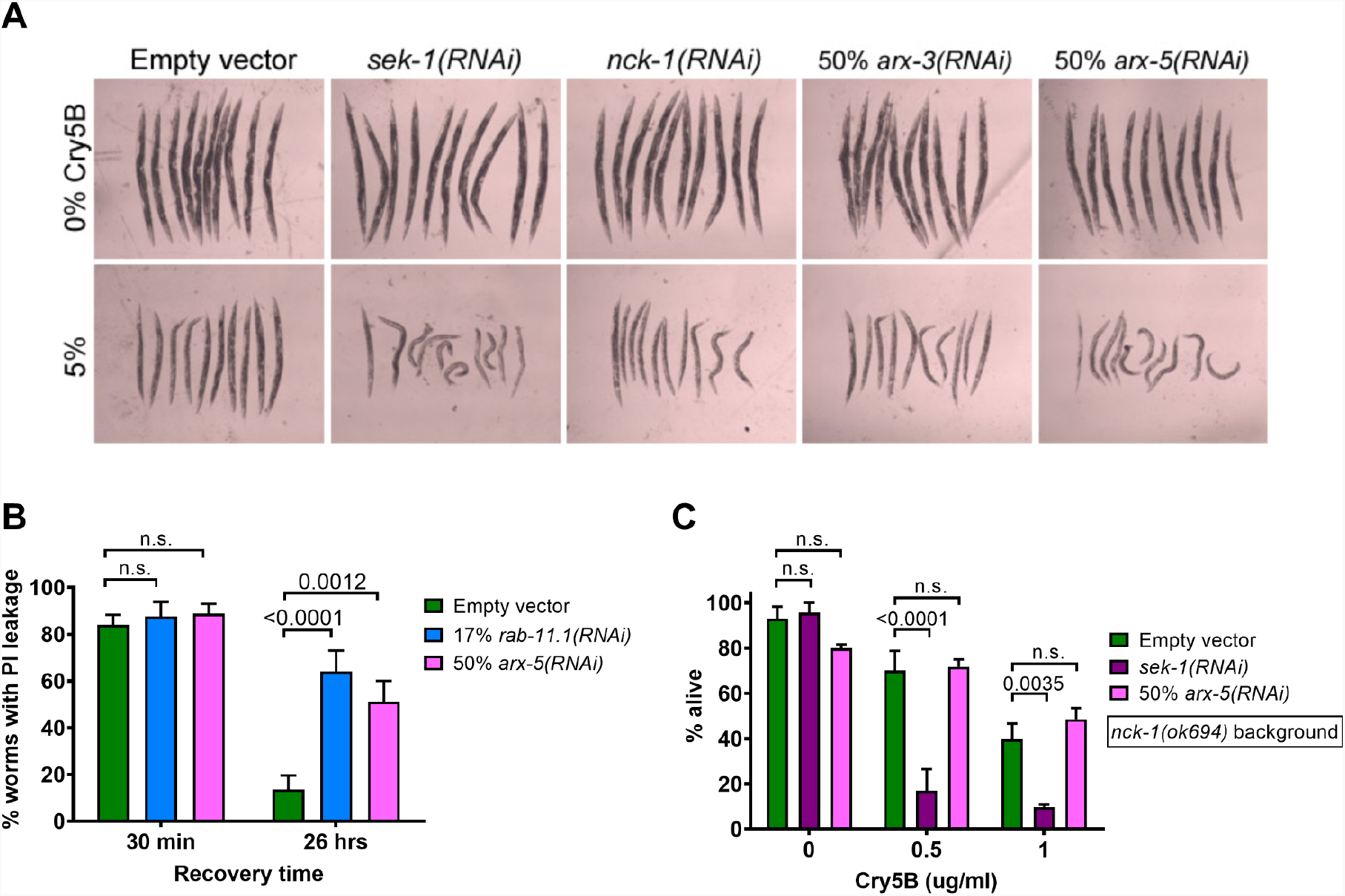
*nck-1*genetically is in the same pathway with the Arp2/3 complex in PFP defense. (A) N2 worms grown on the indicated RNAi bacteria to the L4 stage and then photographed after two days of feeding on the indicated amount of *E. coli-*expressed Cry5B. (B) A pore repair assay identical to the one in Figure 5B, but with N2 worms knocked down for *sek-1* or *arx-5*. (C) Survival of *nck-1(ok694)* worms fed the indicated RNAi bacteria to the L4 stage and then incubated in liquid culture with purified Cry5B for 6 days. The data in Figure 6B represent the mean and SEM of five independent trials, while Figure 6C includes data from three independent trials.

In order to identify additional proteins and processes required for defense against PFPs, we undertook a proteomic approach. *glp-4(bn2)* animals were grown at 25°C to block gonad development and thus remove a significant amount of non-intestinal tissue so that a greater percent of the protein analyzed was derived from the intestine (*glp-4(bn2)* animals have a normal response to Cry5B [14]). L4s were then exposed for 8 hours to *E. coli* either carrying an empty vector or expressing Cry5B. Worms were then processed for proteomic analysis. The list of hits was narrowed down to ones with a significant p-value (p < 0.05), spectral counts > 2, and at least a 2-fold change up or down in expression in Cry5B-exposed worms relative to worms fed with empty vector bacteria. The resulting 386 proteins with increased abundance and 108 proteins with decreased abundance were then analyzed using the PANTHER classification system [38,39]. Analysis of the proteins increased in abundance in the context of Protein Class gave “hydrolase” as the largest category (47 hits). The second largest category was “cytoskeletal protein” (29 hits), and its largest subcategory was “actin family cytoskeletal protein” (21 hits), including 4 subunits of the Arp2/3 complex (ARX1, ARX2, ARX5, ARX7) (Table 1). Furthermore, the largest Molecular Function category in an overrepresentation test of all upregulated proteins was “actin binding” (p = 4.22E-10), with 13 of 55 reference genes represented. This list was mostly a subset of the “actin family cytoskeletal protein” Protein Class and again included the Arp2/3 subunits ARX5 and ARX7. Other noteworthy proteins increased in abundance included components of MAPK signaling, including MEK2, KGB1, and PMK3. Among the proteins with reduced abundance, only one was associated with the Molecular Function term “actin binding”: TWF2, a homolog of the actin-binding protein twinfilin. NCK1 itself was not identified in the proteomic screen.

**Table 1.**
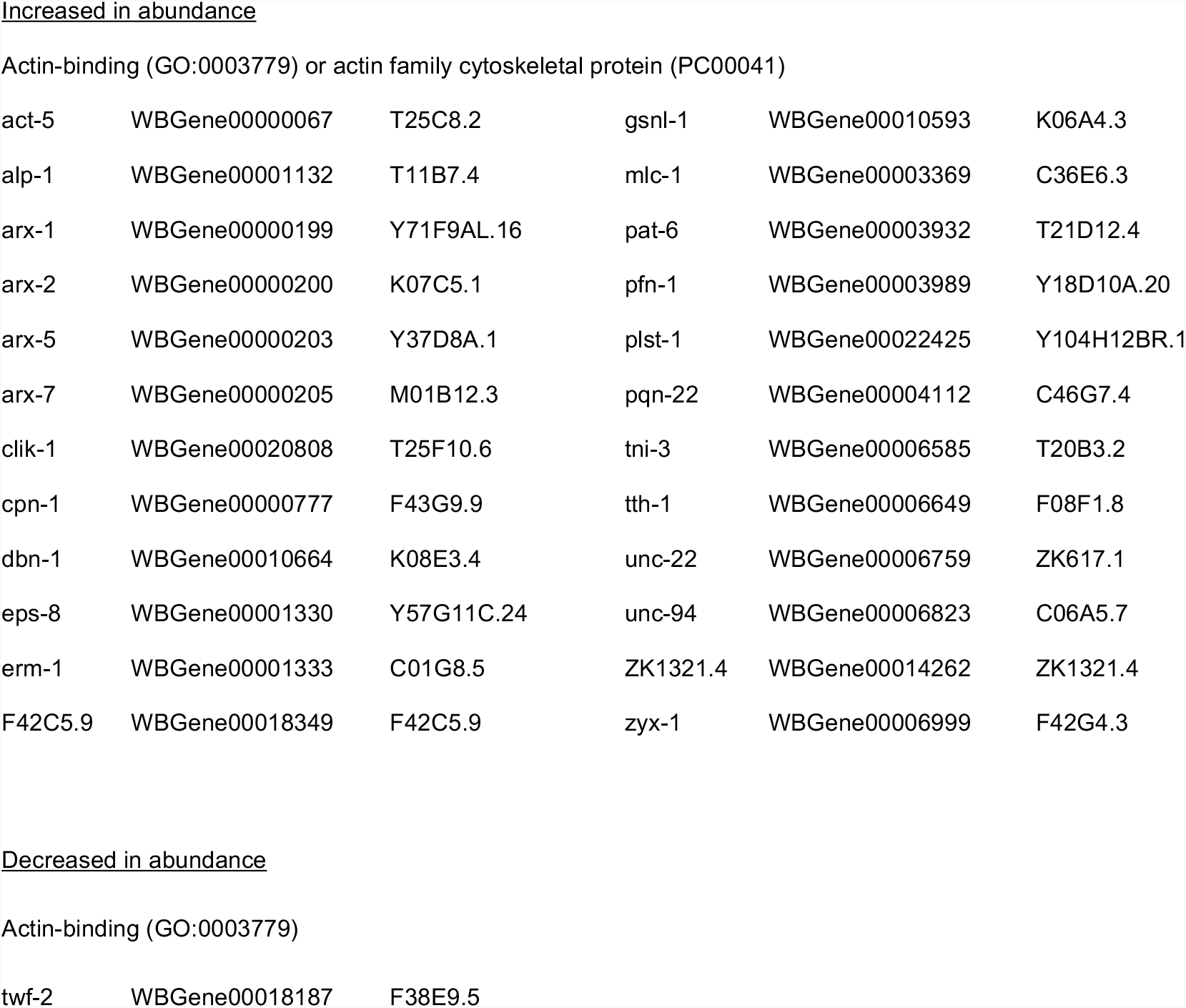
Actin-related *C. elegans glp-4(bn2)* proteins altered by Cry5B feeding as detected by proteomics.

|  |  |  |  |  |  |
| --- | --- | --- | --- | --- | --- |
| act-5 | WBGene00000067 | T25C8.2 | gsnl-1 | WBGene00010593 | K06A4.3 |
| alp-1 | WBGene00001132 | T11B7.4 | mlc-1 | WBGene00003369 | C36E6.3 |
| arx-1 | WBGene00000199 | Y71F9AL.16 | pat-6 | WBGene00003932 | T21D12.4 |
| arx-2 | WBGene00000200 | K07C5.1 | pfn-1 | WBGene00003989 | Y18D10A.20 |
| arx-5 | WBGene00000203 | Y37D8A.1 | plst-1 | WBGene00022425 | Y104H12BR.1 |
| arx-7 | WBGene00000205 | M01B12.3 | pqn-22 | WBGene00004112 | C46G7.4 |
| clik-1 | WBGene00020808 | T25F10.6 | tni-3 | WBGene00006585 | T20B3.2 |
| cpn-1 | WBGene00000777 | F43G9.9 | tth-1 | WBGene00006649 | F08F1.8 |
| dbn-1 | WBGene00010664 | K08E3.4 | unc-22 | WBGene00006759 | ZK617.1 |
| eps-8 | WBGene00001330 | Y57G11C.24 | unc-94 | WBGene00006823 | C06A5.7 |
| erm-1 | WBGene00001333 | C01G8.5 | ZK1321.4 | WBGene00014262 | ZK1321.4 |
| F42C5.9 | WBGene00018349 | F42C5.9 | zyx-1 | WBGene00006999 | F42G4.3 |

|  |  |  |
| --- | --- | --- |
| twf-2 | WBGene00018187 | F38E9.5 |

The proteomics results lent further strength to the hypothesis that actin regulation plays a role in the cellular response to attack by Cry5B. We tested a few of the genes found in the proteomics screen by RNAi knockdown or with mutant strains to see if they sensitized worms to Cry5B. We found that partial knockdown of *erm-1* (Ezrin/Radixin/Moesin-like, involved in linking cortical F-actin to the plasma membrane; [40]) in N2 animals sensitized them to Cry5B feeding (Figure 7A). We obtained the mutant strain *dbn-1(ok925)* (Drebrin-like, an F-actin-binding protein that changes the helical pitch of actin filaments and decreases rate of F-actin depolymerization; [41]), outcrossed it four times, and subjected it to our standard Cry5B LC50 experiment. Consistent with the fact that DBN1 levels were increased in the Cry5B-treated worms in the proteomics experiment, the null mutant strain was hypersensitive to Cry5B relative to N2 (Figure 7B). We also tested for genetic interaction between *erm-1* and either *nck-1*-mediated or *sek-1*-mediated PFP responses. *sek-1(km4)* animals treated with *nck-1* RNAi did significantly shift the IC50 (Figure 7C), in accordance with the earlier genetic interaction data (Figure 2B). As with *nck-1* RNAi, *sek-1(km4)* animals treated with 50% *erm-1* RNAi also showed a significant hypersensitization to Cry5B based on IC50. These data implicate additional actin-interacting proteins, namely Arp2/3, *erm-1* (Ezrin/Radixin/Moesin), and likely *dbn-1* (Drebrin), in the mechanism of an *nck-1*-mediated response to pore-forming proteins that is independent of the p38 MAPK pathway.

**Figure 7.**
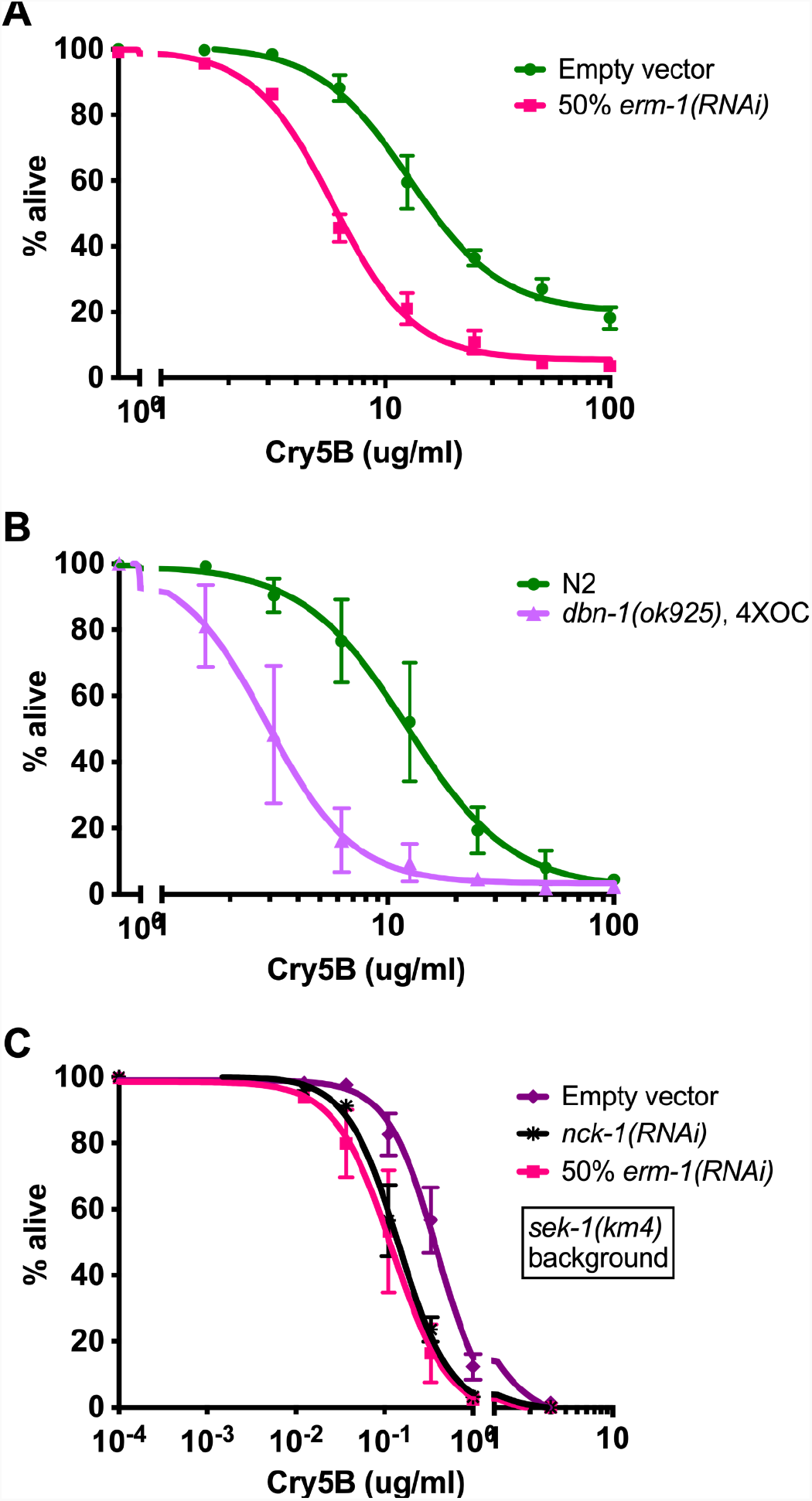
*nck-1* genetically functions in the same pathway as other F-actin-interacting proteins in PFP defense. (A) Wild-type N2 *C. elegans* were grown from the L1 to L4 stage on *E. coli* expressing the indicated RNAi, then transferred to liquid culture with purified Cry5B for 6 days and assayed for survival. (B) N2 or mutant worms were grown on OP50 *E. coli* to the L4 stage, then transferred to liquid culture with purified Cry5B for 6 days and assayed for survival. (C) *sek-1(km4)* mutant *C. elegans* were grown on the indicated RNAi bacteria to the L4 stage and then transferred to liquid culture with varying amounts of purified Cry5B, incubated for 6 days, and scored for survival. All graphs show the mean and SEM from three independent trials.

To support the conclusion that the actin cytoskeleton plays an important role in PFP defenses, we examined whether gross changes in actin abundance occurred in *C. elegans* intestinal tissue upon exposure to Cry5B PFP. We therefore exposed N2 and *nck-1(ok694)* mutant worms to *E. coli*-expressed Cry5B (or empty vector) for 2 hours, then transferred the worms to slides, externalized the intestines, and then stained with FITC-phalloidin to visualize F-actin in the exposed internal tissues. The worms were then analyzed on a confocal scope and images were taken of the stained intestines (Figure 8A). There was an obvious visual decrease in actin staining at the apical surface following Cry5B PFP exposure, and it appeared that *nck-1(ok694)* animals had lower actin staining than wild-type animals in the absence of Cry5B PFP (no toxin) (Figure 8A).

**Figure 8.**
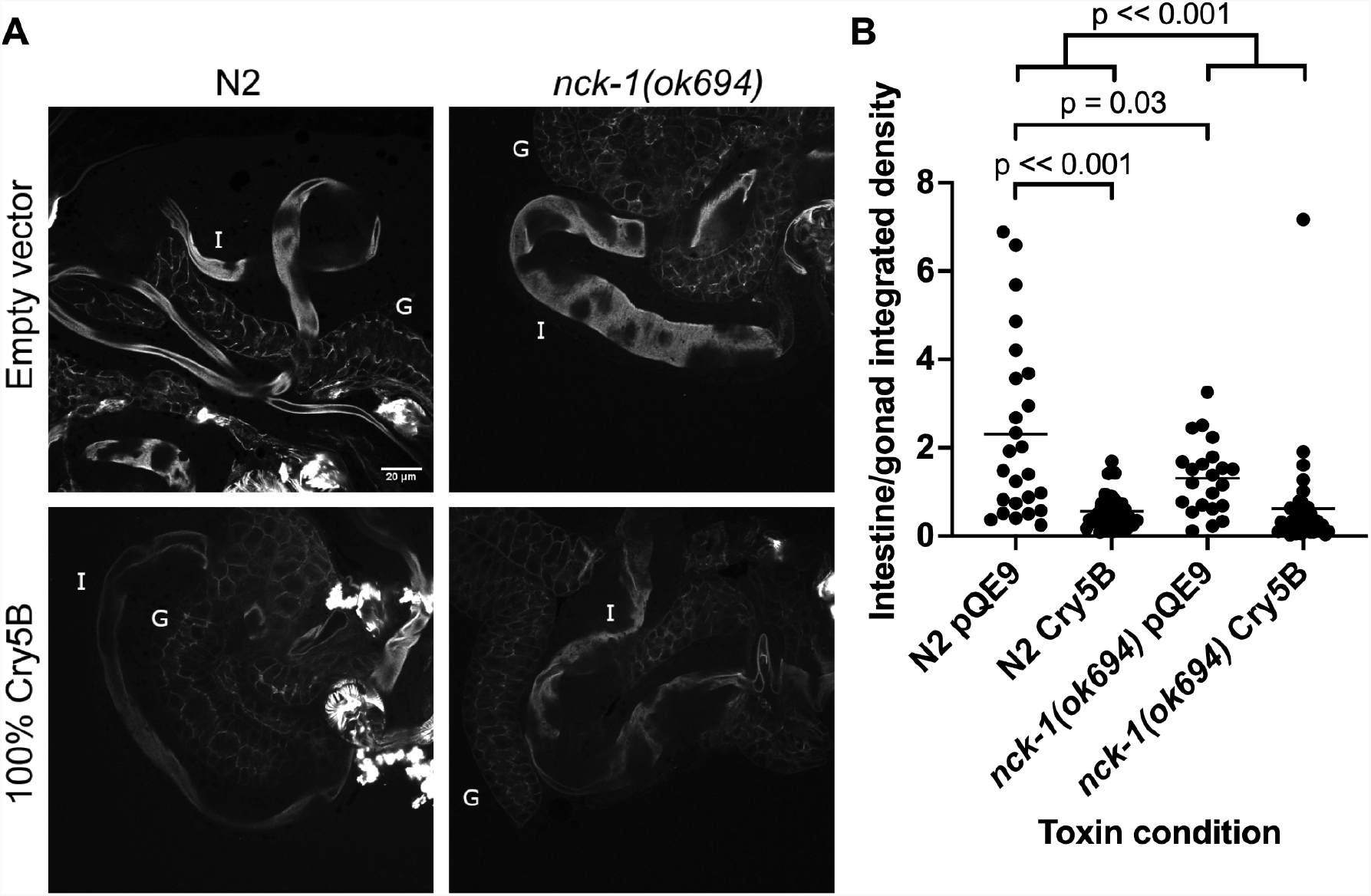
Cry5B exposure causes a reduction in intestinal F-actin levels. (A) Wild-type N2 or *nck-1(ok694) C. elegans* L4 animals were fed *E. coli*-expressed Cry5B or empty vector for 2 hours, permeabilized, and then their internal organs were fixed and stained with FITC-phalloidin. The honeycombed staining is the gonad (labeled G in the upper panels) and the smoother staining is the intestine (I). (B) The integrated density from images (mean pixel intensity x area) of the intestine divided by that of the gonad following the two-hour toxin incubation was graphed. Each dot represents one worm. The data come from three experiments, and the bars show the mean. Statistical analyses are as described in the Methods.

We quantitated the actin staining of the intestine in the images and normalized it to that of the gonad tissue within each photo as we observed that gonadal staining did not seem to fluctuate between PFP-treated and PFP-untreated animals and thus served as an internal PFP-independent control (Figure 8B). When comparing mean F-actin staining levels of N2 hermaphrodites and *nck-1(ok694)* hermaphrodites to each other, *nck-1(ok694)* worms showed a significantly different mean value for intestinal actin staining even before exposure to Cry5B (*i*.*e*., no Cry5B). Following Cry5B exposure, N2 hermaphrodites showed a significant decrease in their intestinal F-actin levels compared to hermaphrodites not exposed to Cry5B. A decrease was also seen in *nck-1(ok694)* animals exposed to Cry5B. However, the magnitude of this decrease was significantly smaller in *nck-1(ok694)* animals than in N2 animals. These data demonstrate that the presence of NCK-1 promotes F-actin polymerization in the intestine and that one of the consequences of intestinal exposure to PFPs is a reduction in F-actin levels.

## Discussion

Here we describe one of the most compelling associations between genes involved in regulating filamentous (F-)actin and *in vivo* intrinsic cellular defenses (INCED) against pore-forming proteins. We find that *nck-1*, a gene known to play a central role in regulated F-actin assembly in cells in response to cell signaling, protects *C. elegans* against small-pore PFPs, genetically functioning in the same pathway as a number of key regulators of F-actin including *C. elegans* Arp2/3 complex (*arx-5*), Ezrin/Radixin/Moesin (*erm-1*), and Drebrin (*dbn-1*). This role in the *nck-1* pathway protecting against small PFP attack is applicable to at least three unrelated PFPs (Cry5B, App6A, VCC) and highly specific in that loss of *nck-1* does not result in increased sensitivity to heat stress, heavy metal stress, high salt, oxidative stress, or pathogenic *P. aeruginosa* bacterial attack. At least part of the role of *nck-1/arx-5* in PFP INCED involves enhancing the ability of *C. elegans* intestinal cell apical membrane to repair the small pores formed by Cry5B.

Specific events that could be tied to F-actin-mediated protection against PFP attack include regulation of clathrin-mediated endocytosis or membrane trafficking (*e*.*g*., in *C. elegans*; [15,42], direct interaction between PFPs and actin [43,44], or generation/maintenance of epithelial junction integrity [45,46].

These results are in contrast to those of the p38 and JNK-like MAPK signaling pathways that are required in *C. elegans* for defense against pathogenic *P. aeruginosa* [27], oxidative stress [28], heat stress [29], and heavy metal stress [14]. This positions *nck-1* as part of a much more specific response to PFPs than any previously characterized.

Consistent with differences in protection against various stressors, *nck-1* and p38 and JNK-like MAPK pathways appear to function differently in PFP defense. Microarrays of *C. elegans* exposed to Cry5B led to the discovery of p38 and JNK-like MAPK signaling as critical mediators of PFP defense, as *C. elegans* that are deficient in either signaling pathway are quite sensitive to small amounts of toxin [13,14]. Indeed, the majority, but not all, of the genes that were upregulated in response to Cry5B exposure became so in a MAPK-dependent manner [13]. Our results here from *hsp-4::gfp* staining and double mutant analyses indicate that *nck-1*/*arx-5*/*erm-1* function separately from the p38 MAPK signaling pathway in PFP INCED. This is the first instance of a major PFP INCED pathway in *C. elegans* functioning independently of a MAPK pathway.

Changes in the actin cytoskeleton by PFPs have been seen in a few instances before, although in general it was not known whether the actin changes are initiated as a means of promoting or resisting pathogenesis [47–49]. An exception was Vega-Cabrera et al. where RNAi of actin led to increased PFP toxicity [44]. Recently, studies on changes in actin remodeling and the role of actin cytoskeleton during plasma membrane damage repair have been reviewed [16]. Interestingly, compounds that stabilize F-actin, like jasplakinolide and phalloidin, were found to hinder the recovery of plasma membrane recovery upon mechanical, laser, or PFP-induced damage. Conversely, F-actin depolymerizing drugs like cytochalasin D and latrunculin increased the speed of repair. There are several important differences between these studies and ours. First, these studies look at repair from membrane damage much larger than that caused by the small-pore PFPs, including by large pore-forming PFPs [5] or by mechanical injury or laser injury. It is possible that defense and repair from smaller pores differs significantly from those caused by much larger plasma membrane disruptions. Second, our study is carried out *in vivo* in an intact epithelium whereas these studies were carried out *in vitro*. In the case of bacterial virulence, this is the actual context in which a PFP attack would occur. Third, it is possible that what is most important is actin dynamics and less so the total amount of F- vs G-actin.

Our data for the first time indicate that an NCK-1 - ARP2/3 - ERM F-actin-regulating pathway is important in PFP INCED in *C. elegans* and, based on the wide range of small-pore PFPs protected against by *nck-1*, likely in other cells as well. Our results support a model in which small-pore PFPs at the *C. elegans* apical intestinal membrane destabilize F-actin, which is counteracted by the activity of NCK-1 - ARP2/3 - ERM, leading to repair of the pores and perhaps other actin-associated INCED processes. Future studies will be directed at determining what other effectors are involved in *nck-1*-mediated PFP defenses and characterizing more precisely how actin dynamics may be involved in INCED against pore-forming proteins.

## Materials and Methods

### *C. elegans* and bacterial strains

*C. elegans* N2 Bristol was maintained using standard techniques [50]. The following strains were used in this study and were purchased from the Caenorhabditis Genetics Center: *glp-4(bn2), glp-4(bn2); rrf-3(pk1426), nck-1(ok694)* following 4X outcross with *lon-2(e678), dbn-1(ok925)* following 4X outcross with *unc-64(e246)*, VP303 *rde-1(ne219);KbIs7[nhx-2P::rde-1], sek-1(km4)*, and SJ4005 *zcIs4[hsp-4::GFP]. E. coli* empty-vector control and Cry5B-expressing strains were JM103-pQE9 and JM103-Cry5B. The App6A gene was subcloned into the *Bam*HI and *Pst*I sites of the vector pQE9 (adding an N-terminal His tag and a seven amino acid extension to the C-terminus) and transformed into JM103 [51]. Feeding RNAi bacterial strains were *Escherichia coli* HT115 with pL4440 vector derived from the Ahringer RNAi library [52] except for rab-11.1 [15]. RNAi clones were confirmed by plasmid DNA sequencing. In all experiments where *arx-5, dbn-1*, or *erm-1* RNAi bacteria were used, a mixture of equal parts HT115 carrying empty pL4440 and or pL4440 bearing the gene-specific RNAi sequence was prepared. In all experiments where *rab-11*.*1* RNAi bacteria was used, a 17% *rab-11*.*1* mixture was prepared in a similar way. *E. coli* OP50, *Pseudomonas aeruginosa* strain PA14 [53], and *Vibrio cholerae* strains CVD109 *Δ(ctxAB zot ace)* and CVD110 *Δ(ctxAB zot ace) hlyA::(ctxB mer) Hg*^*r*^ were used [54].

### Microscopy and Image Editing

Images from qualitative toxicity assays were obtained from assay plates, using an Olympus SZ60 dissecting microscope linked to a Canon Powershot A620 digital camera, and using Canon Remote Capture software or from slides with an Olympus BX60 compound microscope with an UplanFl 10x/0.25NA objective, mounted to a Spot Insight CCD camera and using Spot software. Actin staining images were taken on a Leica TCS SP8 Spectral Confocal DMi8 Inverted Stage Microscope, with LAS X software. All images within an experiment were taken with identical camera settings. Final images were assembled using Adobe Photoshop and/or GIMP.

### Quantitative Cry5B assays in liquid media

Assays for determining LC_50_ of *C. elegans* exposed to purified Cry5B were performed as described [55] with any changes described below. Unless otherwise indicated, all assays were scored after 6 days and included 5-fluoro-2’-deoxyuridine (FUdR; SIGMA) to prevent the appearance of the next generation of larvae that would complicate the experiments [55]. For experiments in which genes were knocked down by feeding RNAi, L1 larvae were synchronized by hypochlorite treatment and plated onto ENG-IA agar plates spread with RNAi bacteria. Animals were grown at 20°C to the L4/young adult stage. OP50 bacteria in the wells were substituted with overnight cultures of RNAi bacteria that were induced with 1 mM IPTG for 1 hr at 37°C before inclusion in wells. Some L1s were plated in parallel on *act-5* or *unc-22* RNAi to verify RNAi effect. In the assays for figure 1B, no FUdR was present and animals were grown to the L4 stage at 25°C. In the assays for figures 1C, 5B, and 5C, *nck-1(ok694)* L4 worms were manually transferred into wells. In the assays for figure 2B and 2C, CuSO_4_ (SIGMA) or NaCl (SIGMA), respectively, were substituted for Cry5B [15]. In the assays for figure 2D, L4 animals were put into wells with only S Media and the indicated concentration of H_2_O_2_ for four hours at 20°C before scoring. In the assays for figures 2B, 2C, and 2D, a parallel qualitative agar plate assay was performed (described below).

### Heat stress assay

VP303 *C. elegans* were synchronized by hypochlorite treatment and grown from the L1 stage on ENG-IA agar plates spread with RNAi bacteria. Some L4 animals were used for a qualitative plate assay (described below). One-day adults were transferred to fresh RNAi plates and placed in a 35°C incubator. At the indicated time-points, plates were removed from the incubator, dead worms (unresponsive to touch by an eyelash pick) were removed from the plate, and the plates were replaced in the incubator.

### *C. elegans* survival assays

PA14 slow-killing assays were performed as described [56], except SK plates seeded with PA14 were incubated overnight at 25C. N2 *C. elegans* were synchronized by hypochlorite treatment and grown from the L1 stage on ENG-IA agar plates spread with RNAi bacteria. Worms were moved to new SK plates at each timepoint.

Some L4 animals were used for a qualitative plate assay (described below). In the *Vibrio cholerae* assays for figures 3B and 3C, CVD109 and CVD110 were inoculated into LB and grown overnight at 30°C. Cultures were then diluted to OD 2.00 and 30 µl were spread on 60mm NG plates and the plates incubated at 25°C overnight. Fifty L4 animals (grown on RNAi bacteria for figure 3B or OP50 bacteria for figure 3C) were transferred to each plate and kept at 25°C. Dead worms were removed at 24 hours and remaining live worms moved to a new set of plates. Final survival counts were taken at 48 hours and those results graphed. Some L4 animals were used for a qualitative plate assay (described below).

### Qualitative agar plate assays

Qualitative agar plate assays were performed as described [55]. *C. elegans* were synchronized and grown to the L4 stage on agar plates spread with either RNAi bacteria or OP50, as required by the experimental design. JM103-pQE9 or JM103-Cry5B bacteria were grown overnight in LB-ampicillin in a 37°C shaker. The next day, the cultures were diluted 10-fold into fresh LB-ampicillin and shaken for 1 hr at 37°C and then 50 µM IPTG was added to induce Cry5B production and the cultures were shaken for 3-4 hours at 30°C. The bacteria were diluted to an OD of 2.00 and spread onto 60mm ENG-IA plates at the indicated concentrations, usually with one set of plates spread with only JM103-pQE9 (0% plates) and another set spread with a 95:5 mixture of JM103-pQE9:JM103-Cry5B (5% plates). Plates were incubated overnight at 25°C and then ten L4 animals were placed on the plates and incubated at 20°C for 2 days. Animals were recovered and placed into glass spot wells with M9 solution with 15 mM sodium azide to paralyze them, and the wells were photographed. In the assays for figure 3A, an additional set of plates were spread with 5% JM103-App6A, prepared identically to JM103-Cry5B.

### Unfolded protein response

Synchronized *hsp-4::gfp* animals were grown to the L4 stage at 20°C on ENG-IA agar plates spread with RNAi bacteria. Animals were then transferred to ENG-IA agar plates that had been spread as described above with either JM103-pQE9 (0% plates) or JM103-Cry5B (100% plates), and incubated at 20°C or 30°C for eight hours. Animals were then washed from the plates and mounted in 0.1% NaN3 in M9, on slides made with 5% agar in water with 0.1% NaN3.

### Pore repair assay

The experiment was performed similarly to previously-described ones [15]. Synchronized L1 animals were grown on RNAi or OP50 at 20°C. L4 stage animals were washed off the plates with ddH2O, rinsed once with ddH2O, then transferred to 100% Cry5B plates and incubated for 1 hr at 20°C. After this Cry5B pulse, worms were washed from the plates with M9 and rinsed once. Then some worms were transferred to fresh RNAi plates or OP50 plates and allowed to recover at 20°C for 26 hr, while others were immediately stained as follows. Worms were incubated in M9 with 5 mg/ml serotonin on a rotator for 15 min at room temperature, after which propidium iodide (Sigma) was added. This was incubated 40-60 min on a room-temperature rotator, after which worms were washed twice with M9 media and mounted on slides. Animals were scored positive for cytosolic PI staining if at least one of the enterocytes in the anterior half of the animal was filled with propidium iodide. At least three independent repeats were performed with 50 animals per treatment.

### Actin staining experiment

N2 or *nck-1(ok694)* L4 animals were transferred to plates that had been spread with JM103 *E. coli* carrying either the pQE9 empty vector or pQE9/Cry5B and incubated at 20C for 2 hrs. Worms were then transferred to polylysine-treated glass slides in a drop of cutting buffer (5% sucrose, 100 mM NaCl, 0.02 mM levamisole (Acros cat. no. 187870100)) and the heads were sliced off using two syringe needles. The worms were fixed in 1.25% paraformaldehyde for 10 min at room temperature then placed in a Coplin jar to wash for 30 min in PBT (1X PBS, 0.1% Tween-20, 0.1% EDTA, 0.05% NaN3). Worms were then treated with 6.6 µM FITC-labeled phalloidin (SIGMA P5282) for 1 hr at room temperature and washed for 1 hr in PBT. Worms were mounted in Vectashield. Image stacks were acquired on a Leica TCS SP8 Spectral Confocal Microscope with DMi8 Inverted Stage Microscope, using LAS X software. Images were analyzed by selecting a middle slice, drawing a region around the intestine and the gonad in ImageJ, and dividing the Integrated Density of the intestine by that of the gonad. In the data set, there were two outlier animals, one in the N2 untreated group and one in the *nck-1(ok694)* untreated group, that showed exceptionally high background (>5X the mean integrated density). These animals were removed from analyses.

### Statistical analyses

Statistical analyses were performed and graphs were generated with GraphPad Prism 9.2.0 software (GraphPad, San Diego, CA). In all figures, p-values are written out, in which “n.s.” denotes “not significant.” Sigmoidal graphed data (Figures 1A, 1B, 1D, 2B, 3B, 3C, 3D, 7A, 7B, 7C) were analyzed using Prism’s nonlinear regression model log(test compound concentration; independent variable) vs. response (% alive; dependent variable) -- variable slope (four parameters). An experimental condition was determined to be significantly different from the control if the 95% confidence intervals of the logIC50 value of the two did not overlap. In bar-graph assays with fewer doses (Figures 1C, 2C, 3A, 4B, 4C, 5B, 6B, 6C), a two-way ANOVA was performed. For Figure 3E, the log-rank test was conducted. In Figure 8B, a two-group comparison was conducted by the Wilcoxon–Mann–Whitney test. The investigated pairs of two-group comparison are: (i) N2 pQE9 versus *nck-1(ok694)* pQE9, (ii) N2 pQE9 versus N2 Cry5B, and (iii) the difference between N2 pQE9 and N2 Cry5B versus the difference between *nck-1(ok694)* pQE9 and *nck-1(ok694)* Cry5B. LC50 and 95% confidence intervals are provided in Table S1.

## Acknowledgments

Some strains were provided by the CGC, which is funded by NIH Office of Research Infrastructure Programs (P40 OD010440).

---

**Table S1.** LC50 values along with 95% confidence intervals associated with experimental results.

|  |  |  |  |  |  |
| --- | --- | --- | --- | --- | --- |
| Fig. no. | Genotype | Treatment | LC50 | 95% CI | 95% CI overlaps with control 95% CI? |
| 1A | N2/L4440(RNAi) | Cry5B | 6.119 µg/mL | 5.392-6.945 | Control |
|  | N2/sek-1(RNAi) |  | 1.217 µg/mL | 1.078-1.328 | No |
|  | N2/nck-1(RNAi) |  | 3.342 µg/mL | 2.691-4.196 | No |
| 1B | *glp-4(bn2); rrf-3(pk1426)/L4440(RNAi)* | Cry5B | 4.37 µg/mL | 3.11-6.818 | Control |
|  | *glp-4(bn2); rrf-3(pk1426)/sek-1(RNAi)* |  | 1.113 µg/mL | ND | - |
|  | *glp-4(bn2); rrf-3(pk1426)/nck-1(RNAi)* |  | 1.791 µg/mL | 1.49-2.114 | No |
| 1D | VP303/L4440(RNAi) | Cry5B | 5.497 µg/mL | 4.465-6.812 | Control |
|  | VP303/sek-1(RNAi) |  | 0.9907 µg/mL | ND | - |
|  | VP303/nck-1(RNAi) |  | 1.76 µg/mL | 1.362-2.189 | No |
| 2B | *sek-1(km4)/L4440(RNAi)* | Cry5B | 0.1385 µg/mL | 0.1184-0.1635 | Control |
|  | *sek-1(km4)/nck-1(RNAi)* |  | 0.04219 µg/mL | 0.02776-0.0689 | No |
|  | *sek-1(km4)/pmk-1(RNAi)* |  | 0.2644 µg/mL | 0.1851-0.4549 | No |
| 3B | VP303/L4440(RNAi) | CuSO<sub>4</sub> | 4.884 mM | 4.373-5.861 | Control |
|  | VP303/nck-1(RNAi) |  | 6.776 mM | 6.024-8.506 | No |
| 3C | N2/L4440(RNAi) | NaCl | 19.51 g/L | 17.42-? | Control |
|  | N2/nck-1(RNAi) |  | 19.4 g/L | 18.54-20.82 | Yes |
| 3D | N2/L4440(RNAi) | H<sub>2</sub>O<sub>2</sub> | 5.179 mM | 4.279-5.635 | Control |
|  | N2/nck-1(RNAi) |  | 5.172 mM | ND | Yes |
| 7A | N2/L4440(RNAi) | Cry5B | 12.78 µg/mL | 10.72-15.64 | Control |
|  | N2/50% erm-1(RNAi) |  | 5.849 µg/mL | 4.944-6.981 | No |
| 7B | N2 | Cry5B | 12.18 µg/mL | 10.43-14.31 | Control |
|  | *dbn-1(ok925)*, 4XOC |  | 2.919 µg/mL | 2.646-3.222 | No |
| 7C | *sek-1(km4)/L4440(RNAi)* | Cry5B | 0.3914 µg/mL | 0.2818-0.6224 | Control |
|  | *sek-1(km4)/50% erm-1(RNAi)* |  | 0.1214 µg/mL | 0.09663-0.1530 | No |
|  | *sek-1(km4)/nck-1(RNAi)* |  | 0.1439 µg/mL | 0.1100-0.1928 | No |

## Notes

### Competing Interest Statement

The authors have declared no competing interest.

